# Endothelial Histone Deacetylase 1 Activity Impairs Kidney Microvascular NO Signaling in Rats fed a High Salt Diet

**DOI:** 10.1101/2023.03.08.531731

**Authors:** Luke S. Dunaway, Anthony K. Cook, Davide Botta, Patrick A. Molina, Livius V. d’Uscio, Zvonimir S. Katusic, David M. Pollock, Edward W. Inscho, Jennifer S. Pollock

**Author notes:** Corresponding Author Jennifer S. Pollock, PhD, Kaul Genetics Building Room 802A, 720 20^th^ Street South, University of Alabama at Birmingham, Birmingham, AL 35233 USA.

## Abstract

**Aim:** We aimed to identify new mechanisms by which a high salt diet (HS) decreases NO production in kidney microvascular endothelial cells. Specifically, we hypothesized HS impairs NO signaling through a histone deacetylase 1 (HDAC1)-dependent mechanism.

**Methods:** Male Sprague Dawley rats were fed normal salt diet (NS; 0.49% NaCl) or high salt diet (4% NaCl) for two weeks. NO signaling was assessed by measuring L-NAME induced vasoconstriction of the afferent arteriole using the blood perfused juxtamedullary nephron (JMN) preparation. In this preparation, kidneys were perfused with blood from a donor rat on a matching or different diet to that of the kidney donor. Kidney endothelial cells were isolated with magnetic activated cell sorting and HDAC1 activity was measured.

**Results:** We found that HS impaired NO signaling in the afferent arteriole. This was restored by inhibition of HDAC1 with MS-275. Consistent with these findings, HDAC1 activity was increased in kidney endothelial cells. We further found the loss of NO to be dependent upon the diet of the blood donor rather than the diet of the kidney donor and the plasma from HS fed rats to be sufficient to induce dysfunction suggesting a humoral factor, we termed <u>P</u>lasma <u>D</u>erived <u>E</u>ndothelial-dysfunction <u>M</u>ediator (PDEM), mediates the endothelial dysfunction. The antioxidants, PEG-SOD and PEG-catalase, as well as the NOS cofactor, tetrahydrobiopterin, restored NO signaling.

**Conclusion:** We conclude that HS activates endothelial HDAC1 through PDEM leading to decreased NO signaling. This study provides novel insights into the molecular mechanisms by which a HS decreases renal microvascular endothelial NO signaling.

## Introduction

Excess salt intake in the form of NaCl is a substantial burden to human health. Globally, it is estimated that elevated sodium consumption accounts for 1.65 million annual deaths from cardiovascular disease.^1^ In the United States, it is estimated that excess salt intake accounts for $18 billion healthcare dollars per year.^2^ A high salt diet (HS) is associated with decreased kidney function, increased cardiovascular disease risk, and increased mortality.^3,4^ While hypertension is one mechanism by which a HS increases mortality, it is not the only mechanism. High salt intake is predictive of cardiovascular deaths even after accounting for blood pressure and other known cardiovascular risk factors.^5^ This may be explained by impaired endothelial nitric oxide (NO) signaling, which occurs in both normotensive and hypertensive subjects on HS. ^6,7^

Endothelial NO, produced by endothelial NO synthase (NOS3), is the major source of NO in the vasculature. NO not only regulates vascular tone as a potent vasodilator but protects against smooth muscle proliferation, endothelial apoptosis, and atherosclerotic plaque formation.^8^ The kidney microvasculature is of particular importance as these arterioles play an essential role in regulating salt and water homeostasis. It has previously been shown that a HS impairs afferent arteriole autoregulatory function.^9,10^ HS impairment of endothelial NO has been reported in multiple vascular beds in both mouse and rat models but, the precise mechanisms by which excess sodium intake mediates loss of NO in the microvasculature remain largely unknown.^11-14^

We have previously shown that a HS increases HDAC1 abundance in the kidney medulla of rats and mice, specifically in the renal tubules.^15,16^ However, the impact of a HS on endothelial HDAC1 is not known. The histone deacetylases (HDACs) are a family of enzymes that deacetylate lysine residues on both histone and non-histone proteins resulting in changes in gene expression and enzyme activity.^17^ The approval of HDAC inhibitors as therapeutics for cancers, epilepsy, and bipolar disorder has sparked interest in repurposing HDAC inhibitors for cardiovascular disease.^18^ HDACs are categorized into four classes. Class I HDACs include HDAC 1-3, and 8; Class II includes HDAC 4-7, 9, and 10; Class III includes Sirtuin 1-7; and Class IV includes HDAC 11.^19^ HDAC1, in particular, regulates both redox homeostasis and NO signaling in the endothelium.^20-22^

Our lab previously reported that *in vitro* HDAC1 over-expression in bovine aortic endothelial cells reduced NO production.^20^ We therefore hypothesized HDAC1 mediates HS induced decreases in NO signaling in the kidney microcirculation. Using the blood perfused juxtamedullary nephron preparation, we investigated the impact of a HS on kidney microvascular NO signaling and its dependence on HDAC1. Our studies indicate that a HS impairs endothelial NO signaling by increasing HDAC1 activity.

## Results

### MS-275 restores vasoconstriction to L-NAME

In order to assess NO signaling in the afferent arteriole, we used the juxtamedullary nephron (JMN) preparation to measure the afferent arteriole diameter in response to increasing concentrations of the NOS inhibitor, L-NAME. As expected, the afferent arteriole from normal salt diet (NS) fed rats constricts in a concentration-dependent manner; however, this constriction is blunted in HS fed rats. Inhibition of HDAC1 with 300 nM MS-275 restores vasoconstriction to L-NAME in HS fed rats (**Figure 1**). MS-275 is a class I HDAC inhibitor which is selective for HDAC1 at 300 nM.^23^ Lower doses of MS-275 (30 nM and 100 nM) did not restore constriction to L-NAME (**Supplemental Figure 1**). Throughout the study, we observed no differences in baseline diameter between groups (**Figures 1, 3 and 4**). In order to assess potential off-target effects of MS-275, we measured the ability of MS-275 to scavenge superoxide using the cytochrome C reduction assay. At the concentrations used in these studies, MS-275 had minimal capacity to scavenge superoxide and did so at a similar level as vehicle (MS-275: 8.07 ± 0.73 % vs Veh: 10.28 ± 0.29 %; *P*=0.49). Additionally, we assessed whether MS-275 or the vehicle, DMSO, constricted the afferent arteriole independent of other stimuli. Afferent arteriole reactivity to DMSO (0.01% vol/vol) was assessed in NS fed rats and reactivity to MS-275 (300 nM) was assessed in HS fed rats. We found that neither of these compounds changed resting baseline diameter (**Supplemental Figure 2a**). Because L-NAME inhibits all three NOS isoforms, we measured the afferent arteriolar diameter in response to increasing concentrations of VNIO (NOS1 specific inhibitor) and 1400W (NOS2 specific inhibitor). NS and HS fed rats displayed similar responses to both of these inhibitors (**Supplemental Figure 2b** and **c**). Additionally, arterioles from HS fed rats did not have an impaired response to the NO donor, sodium nitroprusside (**Supplemental Figure 2d**).

**Figure 1.**
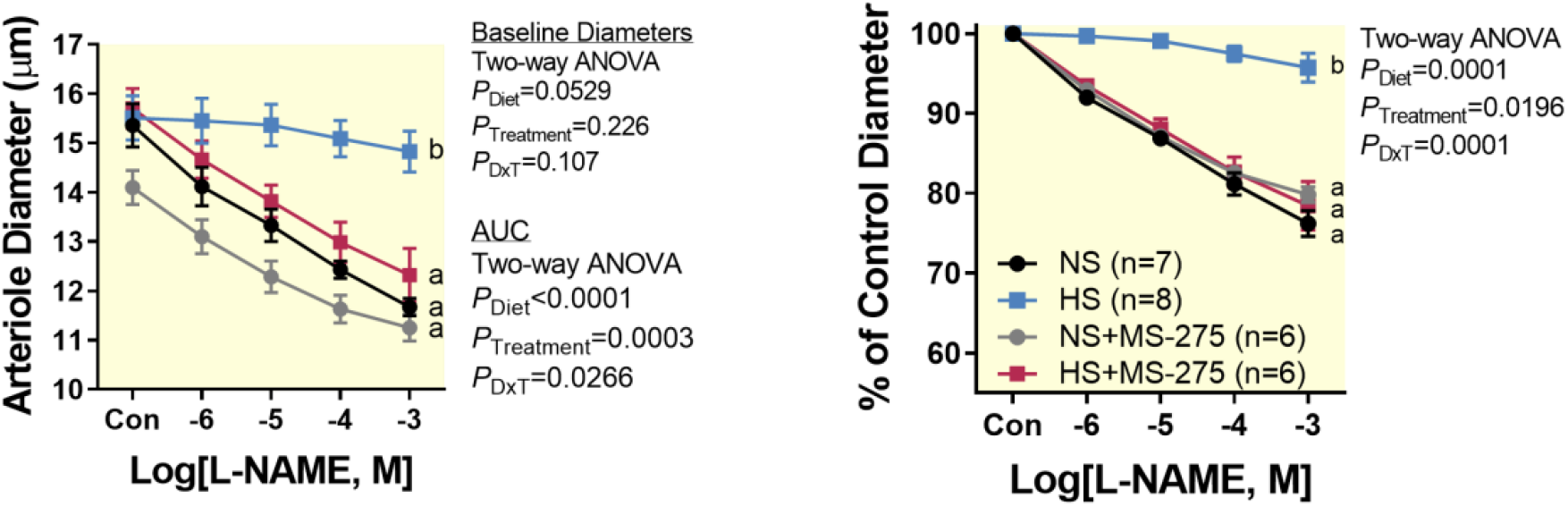
HDAC1 inhibition restores L-NAME constriction in the afferent arterioles of high salt fed rats. L-NAME constricts afferent arterioles from normal salt (NS) fed rats in a concentration-dependent manner. High salt diet (HS) significantly blunts constriction to L-NAME. MS-275 (300 nM) restores constriction to L-NAME in arterioles from (HS). All arterioles were perfused with blood from a rat on a matching diet. Arteriolar diameters are presented as measured (left panel) and normalized to baseline (right panel). Data are plotted as mean±SE. *a* and *b* notate groups with statistically different areas under the curve (AUC). Differences in baseline diameters and AUC were assessed by two-way ANOVA and Holms-Sidak post hoc test.

### Endothelial HDAC1 activity is increased by HS

Using the *in situ* HDAC activity assay, we measured HDAC activity in isolated endothelial cells in the presence of plasma from the same rat. We detected no difference in total HDAC activity but observed increased HDAC1 activity in cells from HS fed rats (**Figure 2a**). When HDAC1 was immunoprecipitated from isolated endothelial cells that were not pretreated with plasma, we measured a decrease in HDAC1 activity with HS (**Figure 2b**). Kidney endothelial cells from NS and HS fed rats had similar levels of class I HDAC abundance as measured by Western blot (**Figure 2c**). We detected similar levels of total HDAC and HDAC1 activity in kidney vessel lysate from NS and HS fed rats (**Figure 2d**). Western blots revealed no difference in class I HDAC abundance in isolated kidney vessels from NS and HS fed rats (**Figure 2e**).

**Figure 2.**
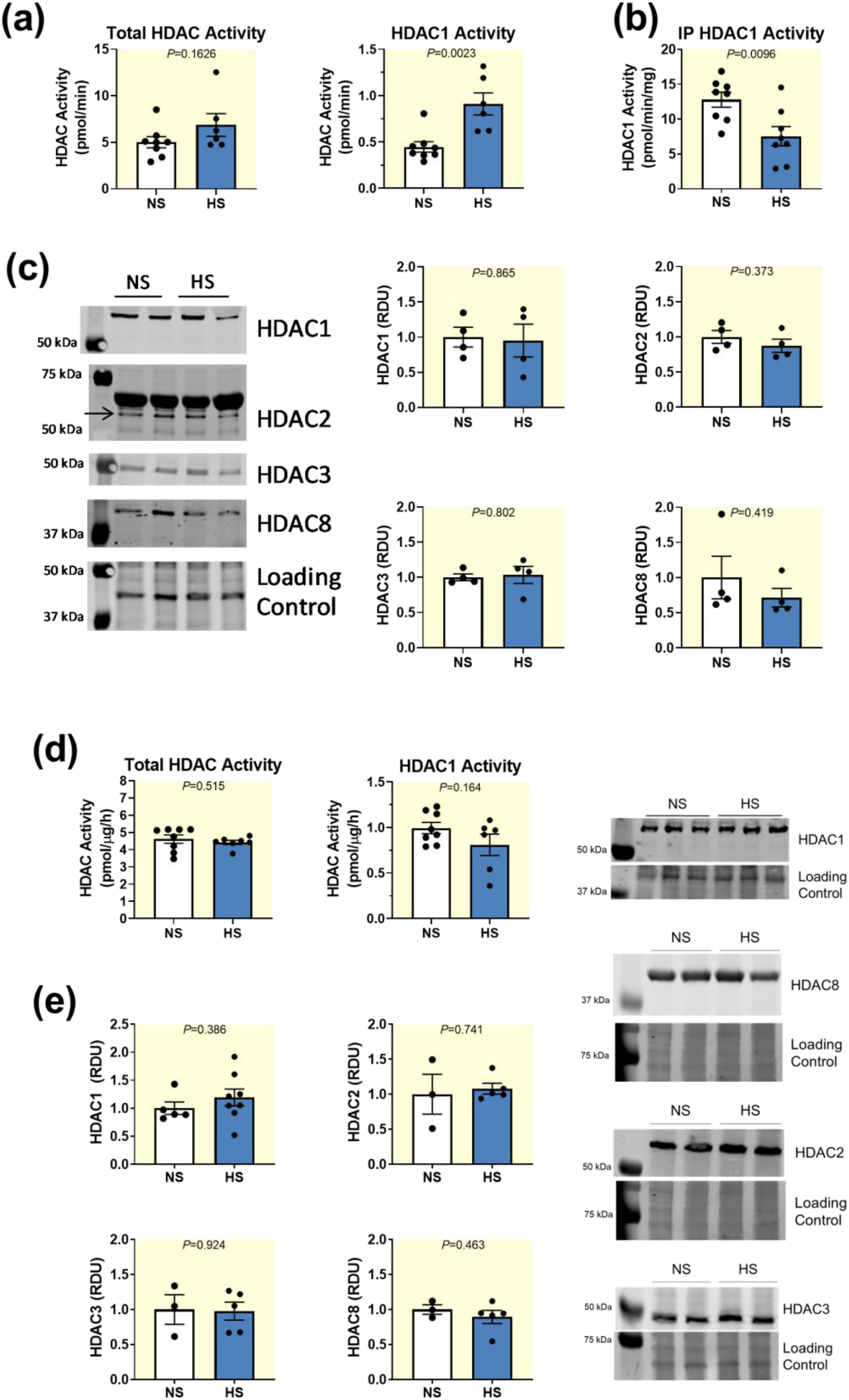
High salt diet increases kidney endothelial HDAC1 activity. (**a**) HDAC1 activity but not total HDAC activity increased with high salt diet (HS) in rat endothelial cells assayed with the *in situ* HDAC activity assay. HDAC1 activity was assessed as the portion of total HDAC activity inhibited by 300 nM MS-275. (**b**) HS decreases the activity of immunoprecipitated HDAC1. HDAC1 was immunoprecipitated from isolated kidney endothelial cells and assayed. (**c**) HS does not change the abundance of class I HDACs in isolated endothelial cells. The arrow notes HDAC2. The large nonspecific band above HDAC2 is BSA which is present in the endothelial cell isolation buffer. REVERT total protein stain was used to assess equal loading. (**d**) HS does not change HDAC activity or HDAC1 activity detected in kidney vessel lysate. HDAC1 activity was assessed as the portion of total HDAC activity inhibited by 300 nM MS-275. (**e**) HS does not change kidney vascular abundance of class I HDACs as measured by Western blot. Coomassie was used to assess equal loading. All data are plotted as mean±SE. Unpaired, 2-tailed Student’s t-test used for all comparisons.

### HS induced disruption of NOS3 is dependent upon a circulating factor

Because HDAC1 activity is increased when assayed in the presence of plasma, we hypothesized that plasma from HS fed rats may be sufficient to impair endothelial NO signaling. In order to test this, we used the JMN preparation to perfuse arterioles from NS fed rats with blood from HS fed rats and vice versa. Arterioles from NS fed rats perfused with blood from HS fed rats had a blunted constriction to L-NAME compared to arterioles from NS fed rats perfused with blood from NS fed rats. Notably, this did not completely abolish constriction to L-NAME as seen in HS arterioles perfused with HS blood. Arterioles from HS fed rats perfused with blood from NS fed rats had similar constriction to L-NAME compared to arterioles from NS fed rats perfused with blood from NS fed rats (**Figure 3a**).

**Figure 3.** Plasma from high salt diet fed rats is sufficient to impair NO signaling. (**A**) Kidneys from normal salt diet (NS) or high salt diet (HS) fed rats were perfused with blood from either NS or HS fed rats. Arterioles perfused with blood from NS fed rats constricted similarly regardless of the diet of the kidney donor. Arterioles from NS fed rats perfused with blood from HS rats had blunted constriction to L-NAME, but not to the level of arterioles from HS rats perfused with blood from HS rats. (**B**) Kidneys from NS fed rats were perfused with reconstituted blood made of erythrocytes from either NS or HS fed rats and plasma from either NS or HS fed rats. Plasma from HS fed rats is sufficient to blunt constriction to L-NAME. (**C**) Arterioles from NS fed rats were perfused with erythrocytes from NS fed rats and plasma from either NS or HS fed rats. 300 nM MS-275 restores constriction to L-NAME in arterioles perfused with plasma from HS fed rats. Arteriolar diameters are presented as measured (left panels) and normalized to baseline (right panels). All data are plotted as mean±SE. *a, b*, and *c* denote groups with statistically different areas under the curve (AUC). Differences in baseline diameters and AUC were assessed by two-way ANOVA and Holms-Sidak post hoc test.

We further investigated whether the HS induced dysfunction was attributed to the plasma or the erythrocyte fractions of the blood. Blood and plasma were prepared as usual (see methods) but rather than recombining erythrocytes with the plasma from the same animal, erythrocytes from HS rats were combined with plasma from NS fed rats and perfused through a kidney from NS fed rats. Likewise, plasma from HS rats were combined with erythrocytes from NS fed rats and perfused through arterioles from NS fed rats. We found that erythrocytes from HS fed rats were insufficient to impair NO signaling, but plasma from HS fed rats blunted L-NAME induced constrictions similar to the levels seen in NS arterioles perfused with both plasma and erythrocytes from HS rats (**Figure 3b**).

We next investigated if plasma from HS rats impaired NO signaling through an HDAC1-dependent mechanism. Indeed, 300 nM MS-275 restored constriction to L-NAME in arterioles from NS rats perfused with plasma from HS rats and erythrocytes from NS fed rats (**Figure 3c**).

### Plasma Components

We performed an initial assessment of several components found in plasma to better understand what may be responsible for the loss of NO signaling (**Table 1**). We investigated several factors that may directly influence NOS3 activity, but found no differences in plasma biopterin, L-arginine, L-ornithine, L-citrulline, arginase activity, and superoxide scavenging capacity. Additionally, we found no differences in plasma electrolytes in our HS model. We further observed that 2 week HS did not increase a select group of cytokines in the plasma.

**Table 1.** Plasma analysis of normal salt (NS) and high salt (HS) fed rats.

|  | **n (NS/HS)** | **NS (mean ± SE)** | **HS (mean ± SE)** | **p-value** |
| --- | --- | --- | --- | --- |
| **<u>Biopterin</u>** |  |  |  |  |
| BH<sub>4</sub> (pmol/mL) | 7/9 | 65.09 ± 11.57 | 53.54 ± 5.59 | 0.311 |
| BH<sub>2</sub> (pmol/mL) | 7/9 | 98.18 ± 7.84 | 87.45 ± 13.28 | 0.530 |
| Total Biopterin (pmol/mL) | 7/9 | 163.3 ± 7.53 | 140.0 ± 17.66 | 0.291 |
| BH<sub>4</sub>:BH<sub>2</sub> | 7/9 | 0.73 ± 0.167 | 0.649 ± 0.0683 | 0.631 |
| **<u>Amino Acid Metabolism</u>** |  |  |  |  |
| L-Arginine (μmol/L) | 8/8 | 162.4 ± 26.56 | 144.1 ± 11.80 | 0.776 |
| L -Ornithine (μmol/L) | 8/8 | 160.5 ± 27.44 | 158.3 ± 16.56 | 0.945 |
| L -Citrulline (μmol/L) | 8/8 | 87.55 ± 8.597 | 100.1 ± 5.814 | 0.247 |
| Arginase Activity (U/L) | 6/5 | 7.96 ± 3.73 | 5.42 ± 1.83 | 0.581 |
| **<u>Antioxidant Capacity</u>** |  |  |  |  |
| Superoxide Scavenging Capacity (%) | 6/5 | 22.04 ± 2.20 | 21.55 ± 1.04 | 0.856 |
| **<u>Electrolytes</u>** |  |  |  |  |
| Na<sup>+</sup> (mmol/L) | 6/7 | 137 ± 0.955 | 136 ± 1.11 | 0.300 |
| K<sup>+</sup> (mmol/L) | 6/7 | 3.4 ± 0.08 | 3.7 ± 0.19 | 0.175 |
| Cl<sup>-</sup> (mmol/L) | 6/7 | 105 ± 1.52 | 104 ± 1.06 | 0.610 |
| HCO<sub>3</sub><sup>-</sup> (mmol/L) | 6/7 | 26.5 ± 0.99 | 27.6 ± 1.37 | 0.539 |
| BUN (mg/dL) | 6/7 | 12 ± 0.83 | 14 ± 0.18 | 0.176 |
| **<u>Cytokines/Chemokines</u>** |  |  |  |  |
| G-CSF (pg/ml) | 10/10 | 61.49 ± 9.763 | 62.02 ± 8.213 | 0.967 |
| Eotaxin (pg/ml) | 10/10 | 24.81 ± 1.623 | 25.3 ± 1.265 | 0.816 |
| IL-1α (pg/ml) | 10/10 | 125.7 ± 19.74 | 129.3 ± 13.76 | 0.883 |
| Leptin (pg/ml) | 10/10 | 5957 ± 589.0 | 4868 ± 318.4 | 0.121 |
| MIP-1α (pg/ml) | 10/10 | 17.27 ± 2.087 | 17.84 ± 1.452 | 0.825 |
| IL-4 (pg/ml) | 10/10 | 176.9 ± 17.30 | 196.1 ± 18.69 | 0.461 |
| IL-1β (pg/ml) | 10/10 | 16.05 ± 2.756 | 16.37 ± 1.919 | 0.296 |
| IL-2 (pg/ml) | 10/10 | 142.4 ± 30.71 | 154.1 ± 21.64 | 0.761 |
| IL-13 (pg/ml) | 10/10 | 36.96 ± 3.978 | 39.47 ± 3.616 | 0.646 |
| IL-10 (pg/ml) | 10/10 | 50.65 ± 6.289 | 59.73 ± 6.130 | 0.315 |
| IL-12 (p70) (pg/ml) | 10/10 | 308.6 ± 50.72 | 334.1 ± 35.25 | 0.685 |
| IFNγ (pg/ml) | 10/10 | 669.4 ± 94.91 | 631.7 ± 55.95 | 0.736 |
| IL-5 (pg/ml) | 10/10 | 234.9 ± 34.16 | 289.0 ± 19.79 | 0.187 |
| IL-17A (pg/ml) | 10/10 | 117.6 ± 20.12 | 122.3 ± 15.06 | 0.854 |
| IL-18 (pg/ml) | 10/10 | 167.5 ± 32.10 | 166.3 ± 19.09 | 0.974 |
| MCP-1 (pg/ml) | 10/10 | 721.9 ± 55.58 | 770.4 ± 50.68 | 0.528 |
| IP-10 (pg/ml) | 10/10 | 78.80 ± 5.487 | 82.66 ± 4.333 | 0.493 |
| VEGF (pg/ml) | 10/10 | 54.82 ± 6.785 | 56.89 ± 4.82 | 0.807 |
| Fractalkine (pg/ml) | 10/10 | 40.86 ± 4.980 | 47.64 ± 6.736 | 0.429 |
| LIX (pg/ml) | 10/10 | 2267 ± 159.7 | 2287 ± 121.8 | 0.920 |
| TNFα (pg/ml) | 10/10 | 9.109 ± 1.379 | 11.17 ± 1.328 | 0.296 |
| RANTES (pg/ml) | 10/10 | 3033 ± 228.1 | 2375 ± 177.8 | 0.0354* |

### Recombinant HDAC1 Activity

Because plasma components may act as direct HDAC inhibitors, we assessed the ability of plasma from NS and HS fed rats to directly inhibit recombinant HDAC1. We found that plasma from HS fed rats inhibited HDAC1 to a significantly greater degree contrary to what we might expect if HDAC1 was directly regulated by plasma components (**Figure 4a**).

**Figure 4.**
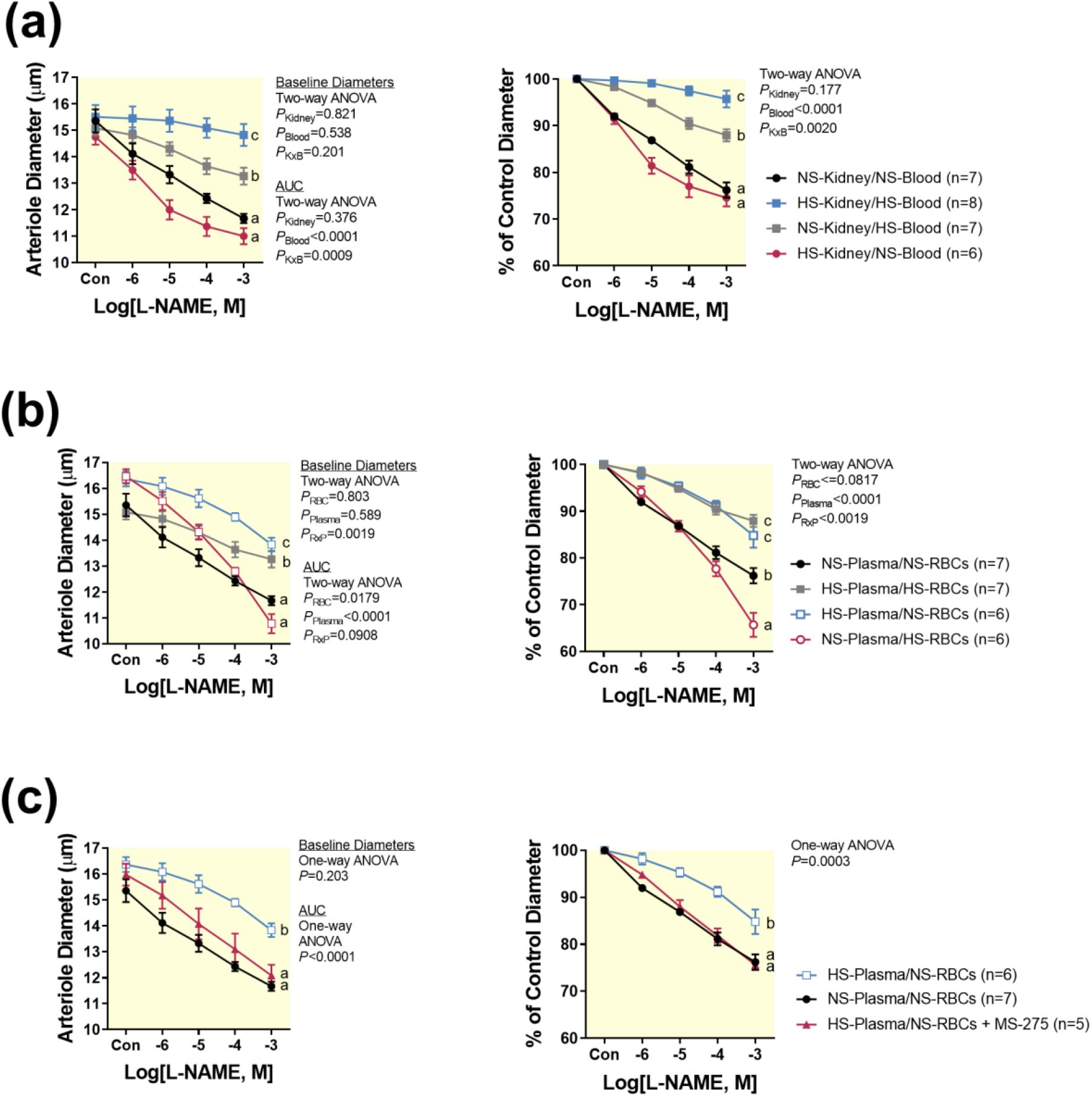
Antioxidants restore L-NAME constriction in the afferent arterioles of high salt fed rats. (A) Recombinant HDAC1 was assayed in the presence of increasing doses of plasma from normal salt (NS) or high salt (HS) fed rats. Differences were assessed with repeated measures two-way ANOVA and Holms-Sidak post hoc test. (B-D) Treatment with 100 U/ml PEG-SOD, 3 µM tetrahydrobiopterin, and 1000 U/ml PEG-catalase restore or partially restore constriction to L-NAME in HS fed rats similar to NS fed rats. Arteriolar diameters are presented as measured (b-d left panels) and normalized to baseline (b-d right panels). All data are plotted as mean±SE. *a, b*, and *c* represent groups with statistically different areas under the curve (AUC). Differences in baseline diameters and AUC were assessed by one-way or two-way ANOVA and Holms-Sidak post hoc test.

### Antioxidants restored constriction to L-NAME

We investigated whether NOS3 phosphorylation or reactive oxygen species (ROS) contributed to the loss of NO signaling seen in HS fed rats. Total NOS3 and phosphorylation of NOS3 at T494, S632, and S1176 were unaffected by HS and HDAC1 inhibition with 300nM MS-275 (**Supplemental Figure 4**). However, pretreatment with PEG-SOD (100 U/ml), the NOS cofactor, BH_4_ (3 µM) restored constriction to L-NAME (**Figure 4b** and **c**). Pretreatment with PEG-catalase (1000 U/ml) largely restored constriction to L-NAME (**Figure 4d)**.

## Discussion

Previous investigations established that a HS impairs endothelial function through a redox-dependent mechanism^6,11^, but this has largely been unexplored in the kidney microvasculature. Renovascular health, in particular afferent arteriole function, is crucial for maintaining kidney function. The afferent arteriole controls glomerular filtration and protects the glomeruli from damaging high pressures. A HS blunts the autoregulatory response in the afferent arteriole, which may contribute to HS-induced kidney damage.^9^ Here we have shown that HS impairs NO signaling in the kidney microvasculature, specifically the afferent arteriole. Furthermore, we found that HS induced changes in plasma composition initiates impaired NO signaling in the afferent arteriole through an HDAC1-dependent mechanism.

NO is produced enzymatically by three enzymes: NOS1, NOS2, and NOS3. There are no known specific NOS3 inhibitors, thus we evaluated arteriolar responses to L-NAME, a pan-NOS isoform inhibitor, as well as NOS1 and NOS2 specific inhibitors. Arterioles from NS and HS fed rats responded similarly to specific inhibition of NOS1 and NOS2 but not to L-NAME. Thus, indicating that NOS3-derived NO signaling is affected by HS in the afferent arterioles. Additionally, HS did not impair the arteriolar response to the NO donor SNP suggesting that HS impairs endothelial specific signaling. Taken together, these findings indicate that HS reduces basal NO production from NOS3 in the kidney microvascular endothelium. This conclusion is consistent with similar findings in other vascular beds,^11-14^ but it should be noted we cannot exclude the possibility that our data may be impacted by unknown potential changes of L-NAME uptake into the cell.

We observed no significant differences in baseline diameter between groups despite the loss of NO signaling with HS. In the microvasculature, NO is only one of many mediators of endothelial-dependent dilation. Other factors such as prostaglandins, epoxyeicosatrienoic acids, or ROS often compensate to regulate basal vascular tone. ^24^ HDAC1 regulates key endothelial functions, including redox homeostasis and NO signaling, through both epigenetic and non-epigenetic mechanisms.^22^ Because acute inhibition of HDAC1 and acute treatment with blood from NS fed rats restored NO signaling, we suggest HDAC1 is acting through an acute, non-epigenetic, signaling mechanism in the setting of HS intake. HDAC1 inhibition had no effect on the NS group suggesting HDAC1 does not suppress basal NO signaling. This is in agreement with previous reports that show increased NO signaling with HDAC1 inhibition or knockdown in pathologic settings, but no further upregulation of NO signaling with HDAC1 inhibition or knockdown in control groups.^25^

We also observed that the blood of HS fed rats was both necessary and sufficient for HS impairment of NO signaling. It should be noted that blood from HS rats did not blunt L-NAME constriction of arterioles from NS fed rats to the same level as HS rats. Because arterioles from HS fed rats retained no dysfunction when perfused with NS blood, we propose the failure to fully blunt L-NAME constriction is due to the acute nature of the exposure to the blood from HS fed rats. Unfortunately, the limitations of this preparation prevent us from testing longer exposures. Furthermore, we observed circulating cells were not necessary for HS impairment of NO signaling, and increased HDAC1 activity was only observed in the presence of plasma from HS fed rats. These findings suggest the presence of a plasma derived endothelial dysfunction mediator (PDEM). When we assayed HDAC1 activity without plasma present, we found HDAC1 activity to be decreased. While we cannot say with certainty what mechanisms are at play, this may suggest activation of negative feedback pathways repressing HDAC1 activity in response to the activation by PDEM. In the absence of plasma, these pathways may result in the apparent reduction in endothelial HDAC1 activity. Based on the sum of the data, we conclude that the plasma fraction harbors factor(s) important in initiating and maintaining kidney microvascular endothelial dysfunction via HDAC1 activation.

We investigated several components of plasma that may have the potential to directly influence NOS3 function. Both NOS3 and arginase compete for the substrate L-arginine and convert it to L-citrulline and L-ornithine respectively. We observed no differences in plasma arginase activity, L-arginine, L-citrulline, or L-ornithine. We also observed that both BH_4_ and PEG-SOD restored NO signaling in arterioles from HS fed rats, but observed no differences in plasma biopterin levels or the ability of the plasma to scavenge superoxide between diets. The effect of HS plasma on NO signaling, is not due to changes in cofactor abundance, competition for substrate, or scavenging of superoxide. These findings are consistent with the conclusion that PDEM works through an HDAC1-dependent mechanism rather than direct modulation of NOS3 enzyme function.

HDAC1 has been shown to regulate both Akt and PKA.^26,27^ Because both of these kinases regulate NOS3 phosphorylation, we investigated the effect of HDAC1 inhibition and HS on NOS3 phosphorylation. HS and HDAC1 inhibition did not affect inhibitory (T494) or activating (S632, S1176) phosphorylation sites. This is in agreement with Guers et al. who found no changes in S1176 phosphorylation in the rat femoral artery in response to HS.^28^ In contrast to our findings, Faraco et al. reported increased T494 phosphorylation in the mouse brain microvasculature in response to HS with 1% saline in the drinking water.^14^ The difference between our findings may be due to differences in the dietary protocol, the species, or vascular bed investigated.

We found the HS-mediated loss of NO signaling to be redox-dependent. Scavenging of either superoxide or hydrogen peroxide completely or partially restored NO signaling. It is well established that ROS impair NO signaling through direct scavenging of NO or oxidation of BH_4_. Because we found BH_4_ restored NO signaling, this may be the mechanism through which oxidative stress impairs NO signaling in our study. Others have shown ROS, and hydrogen peroxide in particular, regulates HDAC1 in endothelial cells^25,29^; therefore, we cannot rule out ROS impairing NO signaling through an HDAC1-dependent mechanism.

HDAC1 activity is modulated by molecules such as short chain fatty acids, inositol phosphates, and intermediates of metabolism.^30-32^ This opens the possibility that PEDM regulates HDAC1 independent of intracellular signaling pathways. We tested the ability of plasma to directly inhibit or activate recombinant HDAC1 and found plasma from HS fed rats shows greater inhibitory capacity than plasma from NS fed rats. This is contrary to what we would expect if the plasma was acting directly on HDAC1, and so it is unlikely that PDEM is directly increasing HDAC1 activity. We propose that the PDEM is activating HDAC1 through an intracellular signaling pathway.

We further assessed plasma components that may have indirect effects on NOS3 function. While some studies suggest changes in plasma sodium mediate endothelial dysfunction on a HS ^33^, we observed no differences in plasma electrolytes in our study after two weeks of HS intake. Others have reported long term HS to increase plasma IL-17A in mice ^14^, but we found two weeks of HS in rats did not increase the abundance of any of the circulating cytokines and chemokines investigated. We did, however, find RANTES to be significantly lower in plasma from HS fed rats compared to NS fed rats. Increased RANTES is associated with endothelial dysfunction and loss of NO in humans and rodent models.^34^ Therefore, we would not expect lower plasma RANTES to be a factor initiating endothelial dysfunction in our model. While we have not yet identified PDEM, we have established the basis for future studies to work towards this end. There are a myriad of humoral factors known to regulate endothelial NO and ROS production such as components of the renin angiotensin aldosterone system, eicosanoids, and microbiome metabolites.^35-37^ Several of these are known to change with dietary interventions and salt in particular. Future studies are in place to identify which, if any, of these are responsible for the HDAC1-dependent loss of NO we observed in this study.

One limitation of our study is the reliance on pharmacologic approaches. To our knowledge, there are no isoform specific substrates for HDACs. Consequently, we had to measure HDAC1 activity as the MS-275 inhibitable portion of HDAC activity in order to measure *in situ* HDAC1 activity. Additionally, we relied on MS-275 to test HDAC1 dependence of HS impaired NO signaling. MS-275 is a class I HDAC inhibitor which has preference for HDAC1 at 300 nM.^23^ Although we saw no antioxidant activity of MS-275, we cannot rule out other off-target effects. This pharmacologic approach is necessary to capture the acute nature of these signaling pathways.

The findings of this study are interesting to consider in light of previous findings from our group regarding the role of HDAC1 in regulating NO signaling in the collecting duct epithelium.^15,16,38-40^ While a HS impairs vascular endothelial NO signaling, it increases tubular NO production. Interestingly, both of these mechanisms are dependent upon HDAC1. It remains unknown how HDAC1 mediates seemingly opposite effects in these two cell types.

Based on the results of this study, we propose the following model shown in **Figure 5**. We put forward that HS intake results in the presence or increase of PDEM which increases HDAC1 activity through an intracellular signaling pathway. Thus, kidney microvascular NO signaling is impaired by an HDAC1 and ROS-dependent mechanism. We further speculate that this mechanism is most likely present in other microvascular beds that are known to have HS-mediated loss of NO signaling.

**Figure 5.**
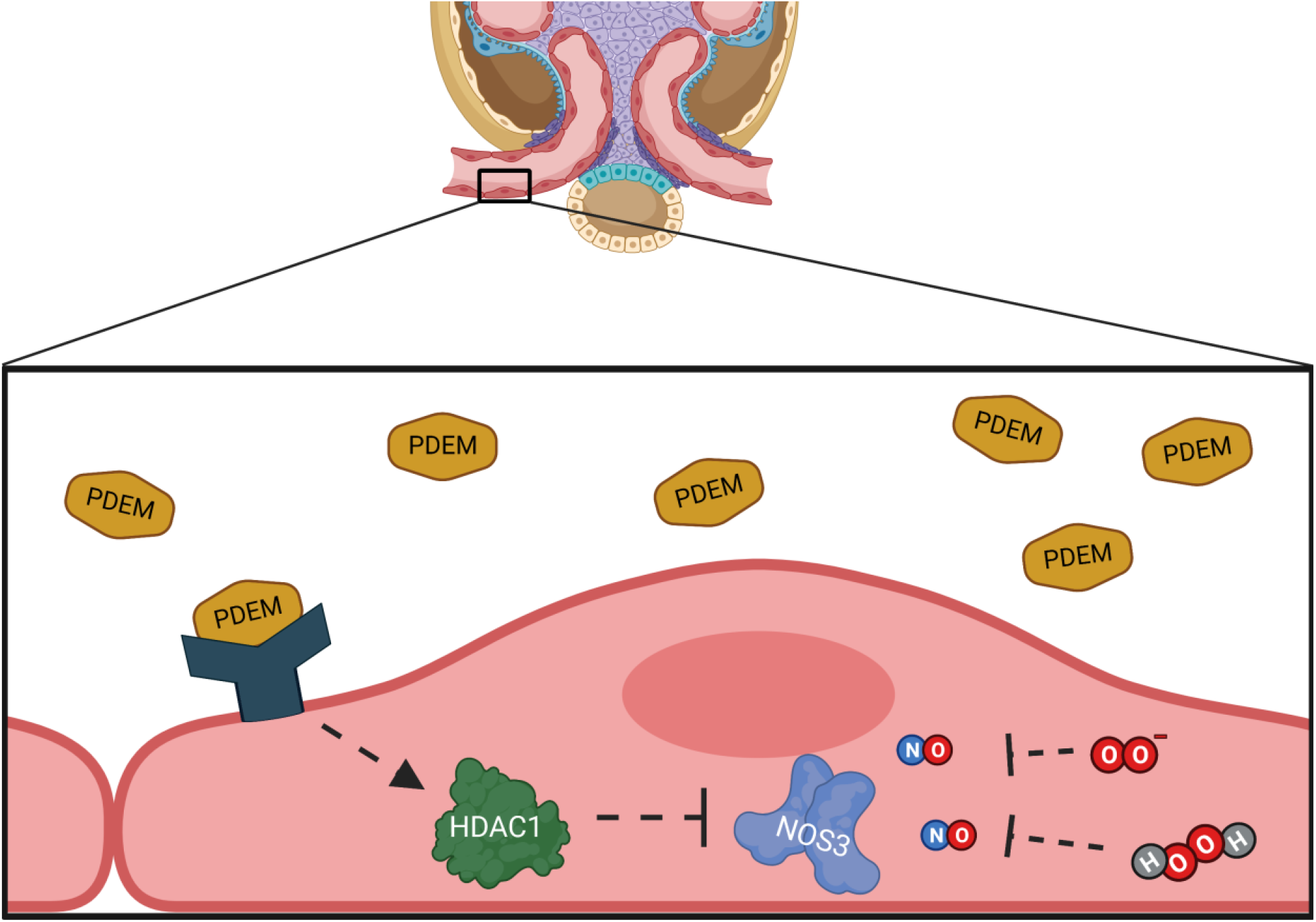
Hypothetical Model. High salt diet promotes the presence of a plasma derived endothelial dysfunction mediator (PDEM) which activates kidney endothelial HDAC1 through an intracellular signaling mechanism. Both increased HDAC1 activity and reactive oxygen species (O_2_^-^ and H_2_O_2_) contribute to the loss of NO signaling in the juxtamedullary afferent arteriole. Created with BioRender.com.

### Perspectives

Loss of endothelial NO precedes the onset of cardiovascular disease, but assessing endothelial NO is not practical in clinical settings. Here we have uncovered evidence of a humoral factor, PDEM, that disrupts endothelial NO signaling in HS fed rats and have paved the way for future work to identify this factor. The identification of PDEM may provide a novel biomarker that is mechanistically linked to loss of NO signaling in the microvasculature allowing for early detection of cardiovascular risk and implementation of disease prevention strategies.

Several studies have shown HS impairs NO signaling through a ROS-dependent mechanism, but the pathways involved have remained elusive. Furthermore, antioxidant therapies have been largely unsuccessful in clinical studies.^41^ HDAC1 has been shown to repress NO signaling and promote oxidative stress in multiple disease settings.^21,25,29^ Here we expand these studies showing increased HDAC1 activity represses NO signaling in HS fed rats. Four HDAC inhibitors are FDA approved and isoform specific inhibitors are undergoing clinical trials. Of relevance to this study, MS-275 has been in multiple phase I clinical trials and has shown promising tolerability.^42-45^ Utilization of HDAC inhibitors to restore NO signaling in cardiovascular disease warrants further investigation.

## Methods

### Animal Model

Male Sprague Dawley rats were purchased from Envigo at 225 - 249 g (Indianapolis, IL) and were kept in a 12-hour lights on and off schedule with lights turning on at 7 am. The rats were fed *ad libitum* and randomly assigned to either a normal salt NIH-31 diet (NS; 0.49% NaCl) or matching high salt (HS; 4% NaCl) (Envigo, TD.150756 or Envigo, TD.92012) diet for 13-15 days. Rats had access to tap water *ad libitum*. The rats were euthanized under isoflurane for tissue and plasma collection between 7 am and 10 am. Blood was collected with heparin and centrifuged at 4000 rpm for 15 minutes (Sorvall Legend X1R Centrifuge, Thermo Scientific). Plasma was filtered with 0.22 µm filters. All animal care and procedures were approved by the Institutional Animal Care and Use Committee of the University of Alabama at Birmingham and in accordance with the National Institutes of Health Guide for the Care and Use of Laboratory Animals.

### Blood Perfused Juxtamedullary Nephron Preparation

The blood perfused juxtamedullary nephron preparation was performed as before. ^46^ In brief, rats were anesthetized with pentobarbital (50mg/kg intraperitoneally) or Inactin (100 mg/kg intraperitoneally). Blood was collected into a heparinized syringe. Blood was pooled from two rats for one experiment. After centrifugation, the plasma fraction was collected and filtered through a 0.45 um and 0.2 um filter. The buffy coat was discarded and erythrocytes were washed twice with 0.9% saline. The plasma and erythrocytes were recombined with a hematocrit of approximately 33% and passed through a 5 µm nylon mesh. The right kidney of one rat was dissected out, cannulated, and perfused with Tyrode’s solution containing 6% BSA (Calbiochem, La Jolla, CA). After dissection and visualization of the afferent arteriole, the kidney was perfused with the reconstituted blood and investigated on a microscope stage at 37°C using videomicroscopy.

Where indicated, reconstituted blood from NS fed rats were perfused through the kidneys of HS fed rats and vice versa. In order to separate the effect of the erythrocyte and plasma fractions, the washed erythrocyte fraction from NS fed rats was recombined with the filtered plasma fraction from HS fed rats and vice versa. These were then perfused through a kidney taken from a NS fed rat. The afferent arteriole diameter was measured in response to Nω-Nitro-L-arginine methyl ester hydrochloride (L-NAME), 1400W, or vinyl-L-NIO hydrochloride (VNIO) at increasing concentrations. These nitric oxide synthase inhibitors were administered via the 1% albumin superfusate solution. The arteriole was exposed to each inhibitor concentration for 5 minutes. Arteriole diameter was measured at a fixed location at 12 second intervals. The diameters in the last 2 minutes of each 5-minute interval were averaged. The percent of control diameter was calculated by dividing the diameter at each concentration of L-NAME by baseline diameter. PEG-SOD (100 U/ml) or PEG-catalase (1000 U/ml) (Sigma-Aldrich, St. Louis, MO) were administered in the blood perfusate which provides about 30 minutes of equilibration and remains throughout the experiment. 3 µM tetrahydrobiopterin (Schircks Laboratories, Switzerland) and 300 nM MS-275 (Cayman Chemical, Ann Arbor, MI) were administered in 6% BSA perfusate during dissection which lasts approximately 1 hour and the blood perfusate for the duration of the experiment.

### Endothelial Cell Isolation

Single cell suspensions were generated as previously described.^47^ After blood collection, the rats were perfused with ice cold Ca^2+^ and Mg^2+^ free PBS to reduce the number of erythrocytes in the final cell suspension. After decapsulation, kidneys were minced. Both kidneys were incubated in 10 ml enzymatic solution (1 mg/ml collagenase A (Roche Diagnostics, Germany), 2.5 mg/ml trypsin (Sigma-Aldrich, St. Louis, MO), 0.2 mg/ml DNAse I (Roche Diagnostics, Germany)) for 15 minutes shaking at 80 RPM at 37°C. The tissue was further mechanically disassociated by pipetting up and down. After allowing the dissociated tissue to settle, the supernatant was removed and placed on ice. 10 ml of enzymatic solution was added to the remaining tissue and incubated at 37°C for 15 minutes shaking at 80 RPM. This was repeated once more for a total of three incubation periods. The collected cell suspension was centrifuged for 4 minutes at 800x*g* at 4°C. The pellet was resuspended in 10 ml buffer 1 (NaCl 130 mM, KCl, 5 mM, CaCl_2_ 2 mM, glucose 10 mM, sucrose, 20 mM, HEPES 10 mM) and filtered through a 70 µm filter. The filter was washed with 15 ml of buffer 1. The filtrate was then passed through a 40 µm filter. The filter was washed with 25 ml of buffer 1. The filtrate was then centrifuged for 4 minutes at 800x*g* at 4°C. Cells were then suspended in 0.5% bovine serum albumin in phosphate buffered saline (PBS) pH 7.2. Endothelial cell isolation kit for rat (Miltenyi Biotec Inc., Auburn, CA) was used to isolate endothelial cells from this cell suspension following manufacturer protocol. Isolated endothelial cells were flash frozen and stored at -80°C after isolation or immediately assayed as described below for specific assays. Enrichment of endothelial cells was verified by assessing NOS3 abundance by Western blot (**Supplemental Figure 3**).

### In Situ HDAC Activity Assay

Total HDAC activity was assessed with the in situ HDAC activity fluorometric assay kit (Sigma Aldrich, St. Louis, MO). After isolation, kidney endothelial cells were incubated in plasma from the same donor rat for one hour in a 37°C water bath. Cells were then centrifuged 1000x*g* for 5 minutes at room temperature and resuspended in the reaction mix (89% HBSS, 10% plasma, 1% substrate). Cells were plated at 50,000 cells/well in a 96 well plate. The reaction proceeded for 2 hours at 37°C in presence of either 300 nM MS-275 (Cayman Chemical, Ann Arbor, MI) or vehicle (0.01% DMSO). After 2 hours, the deacetylated substrate was developed following manufacturer instructions. Each condition was assayed in duplicate. The portion of activity inhibited by MS-275 was attributed to HDAC1. MS-275 is a class I HDAC inhibitor with an IC_50_ of 300 nM and 8 µM for HDAC1 and HDAC8 respectively.^23^ Signal from background wells containing only the reaction mix was subtracted from each sample. The amount of generated fluorescent product was calculated using a standard curve generated with the provided standards and divided by 120 minutes to calculate the activity in pmol/min. Vehicle treated samples were considered total HDAC activity. The activity of MS-275 treated samples was subtracted from those treated with vehicle to calculate HDAC1 activity.

### Immunoprecipitated HDAC1 Activity

HDAC1 activity was measured using the HDAC1 immunoprecipitation & activity assay kit (BioVision, Milpitas, CA) with isolated endothelial cells. Frozen endothelial cells were sonicated 10 times on setting 3 (Sonic Dismembrator Model 100, Fisher Scientific) in the proprietary lysis buffer provided with protease inhibitors (10 µM leupeptin, 2 µM pepstatin A, 0.1% aprotinin, 0.35 mg/ml PMSF) and phosphatase inhibitor cocktail A (Santa Cruz, Dallas, TX). Protein was quantified using the Bradford assay (Bio-Rad, Hercules, CA). Assay was performed per manufacturer protocol. Signal from the IgG negative control was subtracted from each sample. The amount of fluorescent product produced was calculated using a standard curve generated with the provided standards and divided by the reaction time (120 minutes) and assayed lysate protein amount (0.075 mg) to calculate the activity in pmol/min/mg.

### Whole kidney vessel isolation

Kidneys were dissected out of the rats and decapsulated. The kidneys were placed in a dish of ice cold physiologic saline solution (130 mM NaCl, 14.9 mM NaHCO_3_, 5.5 mM glucose, 4.7 mM KCl, 2.4 mM MgSO_4_, 2.16 mM CaCl_2_, 1.2 mM KH_2_PO_4_, 20 µM EDTA) between two 70 µm microsieves (BioDesign Inc. of New York, New York, NY). Kidney vessels were dissected with gentle grating for about 2 minutes. To assess NOS3 phosphorylation, vessels were incubated in physiologic saline solution for 1 hour with either 300 nM MS-275 or vehicle (0.01% DMSO) then snap frozen in liquid nitrogen. This technique favors the isolation of larger preglomerular arteries.

### Kidney Vascular HDAC Activity

Isolated kidney vessels were homogenized in lysis buffer (20 mM Tris, 150 mM NaCl, 0.5% Triton X-100, 1 mM EDTA, 1 mM EGTA, pH 7.5) with protease inhibitors (10 µM leupeptin, 2 µM pepstatin A, 0.1% aprotinin, 0.35 mg/ml PMSF) and phosphatase inhibitor cocktail A (Santa Cruz, Dallas, TX) using a dounce homogenizer then sonicated 10 times on setting 7 (Sonic Dismembrator Model 100, Fisher Scientific). The homogenate was centrifuged at 1000x*g* for 5 minutes at 4°C. Protein was quantified by Bradford assay (Bio-Rad, Hercules, CA). 35 ug of protein was assayed for HDAC activity (HDAC activity fluorometric assay kit (BioVision Inc, Milpitas, CA) in the presence of either 300 nM MS-275 (Cayman Chemical, Ann Arbor, MI) or vehicle (0.01% DMSO). Each condition was assayed in duplicate. The portion of activity inhibited by MS-275 was attributed to HDAC1. Signal from the trichosatin A (TSA) treated cells was subtracted as background from each sample. The amount of fluorescent product obtained was calculated using a standard curve generated with the provided standards and divided by the reaction time (0.5 hours) and assayed protein amount (35 µg) to calculate the activity in pmol/µg/hour. Vehicle treated samples were considered total HDAC activity. The activity of MS-275 treated samples was subtracted from those treated with vehicle to calculate HDAC1 specific activity.

### Western Blot

Endothelial cells were lysed immediately following isolation for Western blotting analysis. Endothelial cells were sonicated 10 times on setting 3 (Sonic Dismembrator Model 100, Fisher Scientific) in lysis buffer (20 mM Tris, 150 mM NaCl, 0.5% Triton X-100, 1 mM EDTA, 1 mM EGTA, pH 7.5) with Halt Protease Inhibitor Cocktail including EDTA (Thermo Scientific) and 0.35 mg/ml PMSF and phosphatase inhibitor cocktail A (Santa Cruz, Dallas, TX).

Frozen kidney vessels were homogenized on a Dounce homogenizer in vessel lysis buffer (50 mM Tris, 0.1 mM EDTA, 0.1 mM EGTA, 250 mM sucrose, 10% glycerol, 0.5% triton X-100, 0.1% SDS, 0.5% sodium deoxycholate, 0.1% BME, pH 7.4) with protease inhibitors (10 µM leupeptin, 2 µM pepstatin A, 0.1% aprotinin, 0.35 mg/ml PMSF) and phosphatase inhibitor cocktail A (Santa Cruz, Dallas, TX). Homogenate was flash frozen then thawed on ice. The homogenate was then sonicated 10 times on setting 7 (Sonic Dismembrator Model 100, Fisher Scientific) and incubated at 4°C while rocking for 30 minutes. Lysate was centrifuged at 1000x*g* for 5 minutes at 4°C.

For both kidney vessels and endothelial cells, total protein was quantified with a Bradford assay (Bio-Rad, Hercules, CA). Kidney vessel lysates (40 µg) and endothelial cell lysates (10 µg) were electrophoresed on 8% SDS-PAGE gels and transferred to PVDF membranes. Membranes were stained with antibodies against HDAC1 (1:1000, ab109411, abcam, Cambridge, MA), HDAC2 (1:1000, ab32117, abcam, Cambridge, MA), HDAC3 (1:1000, ab32369, abcam, Cambridge, MA), HDAC8 (1:500 for kidney vessels and 1:1000 for endothelial cells, ab187139, abcam, Cambridge, MA), NOS3 (1:1000, 610297, BD Biosciences, location), phospho-S632 NOS3 (1:200, 07-562, Upstate, Lake Placid, NY), phospho-T494 (1: 200, 9574S, Cell Signaling Technology, Danvers, MA), or phospho-S1176 (1:500, 9571L, Cell Signaling Technology, Danvers, MA) followed by incubation with goat anti-mouse or goat anti-rabbit secondary antibody (1:1000; Invitrogen). Loading was assessed by REVERT Total Protein Stain (LI-COR Biosciences) following manufacturer protocol or Coomassie where indicated. For Coomassie staining, blots were incubated in TBST for 5 minutes shaking at room temperature. Blots were incubated in stain solution (1.5 mg/ml Coomassie Brilliant Blue R-250 Dye (Fisher), 0.5% vol/vol acetic acid, 25% vol/vol MeOH, 74.5% vol/vol H_2_O) for 5 minutes shaking at room temperature. After briefly rinsing in water, the blots were incubated in destain solution 1 (0.5% vol/vol acetic acid, 25% vol/vol MeOH, 74.5% vol/vol H_2_O) for 5 minutes shaking at room temperature. The blots were rinsed in destain solution 2 (1% vol/vol acetic acid, 81% vol/vol MeOH, 18% vol/vol H_2_O) briefly agitating by hand. Finally, the blots were rinsed in water and left to dry. All blots were imaged on Odyssey CLx Infrared Imaging System (LI-COR Biosciences). Densitometry was quantified with LI-COR Image Studio Lite. Expression of each sample was first normalized to either REVERT or Coomassie and presented relative to NS fed rats.

### Plasma Analysis

Plasma biopterin was measured as described.^48^ Plasma L-arginine, L-ornithine, and L-citrulline were quantified with HPLC-MS as previously described.^49^ Arginase activity was assessed with an arginase activity assay kit (Sigma Aldrich, St. Louis, MO) following manufacturer instructions. Activity is reported in U/L where one unit is equal to 1 micromole of product generated per minute.

In order to assess the ability of plasma and MS-275 to scavenge superoxide, we used a xanthine oxidase based cytochrome C reduction assay reported in detail by Quick et al.^50^ These values are presented as percent reduction in cytochrome C oxidation (%).

Plasma electrolytes and blood urea nitrogen were measured with iSTAT EC8+ cartridges (Abbott Laboratories, Princeton, NJ) per manufacturer protocols.

Cytokine and chemokine levels were quantified in serum with the rat-specific, premixed, 27-plex Milliplex® panel kit RECYTMAG65K27PMX (Millipore-Sigma) and using the MagPix® instrument platform with xPONENT® software (Luminex Corp). The readouts were analyzed with the standard version of Milliplex® Analyst software (Millipore-Sigma and Vigene Tech). Some cytokines/chemokines that were below the detectable range were excluded.

### Recombinant HDAC1 Activity Assay

The ability of plasma to inhibit recombinant HDAC1 was assessed with HDAC1 inhibitor screening assay kit (Cayman Chemical, Ann Arbor, MI) following manufacturer instructions. Plasma was tested at 10 µl, 20 µl, and 40 µl in a total reaction volume of 170 µl. Each reaction included 200 µM substrate (K_m_ = 100 µM). The data are presented as percent initial activity. The signal from wells lacking HDAC1 were defined as the background signal and subtracted from the signal of each sample. This was divided by the activity of untreated HDAC1 to calculate the percent of initial activity.

### Statistical Analysis

Statistical analysis was performed with GraphPad Prism 8. Data are plotted as mean±SE. Unpaired, two-tailed Student’s t-test, one-way ANOVA, two-way ANOVA, and two-way ANOVA repeated measures were used as appropriate and indicated in figure legends. Holms-Sidak was used for post hoc analysis. *P*<0.05 was considered statistically significant. For multiple comparisons *a, b*, and *c* denote significantly different groups such that groups marked with the same letter are not significantly different.

## Acknowledgements

This work was supported by National Institutes of Health (NIH) F31 HL149235 and T32 GM109780 to LSD; F31 HL151264 to PAM; DK044628 to EWI; P01 HL136267 to JSP, DMP, and EWI; and by the American Heart Association 15SFRN24450002 to JSP, DMP, and EWI.

## Data Availability Statement

The data that support the findings of this study are available from the corresponding author upon reasonable request.

