## Supplemental Figures for "Endothelial Histone Deacetylase 1 Activity Impairs Kidney Microvascular NO Signaling in Rats fed a High Salt Diet"

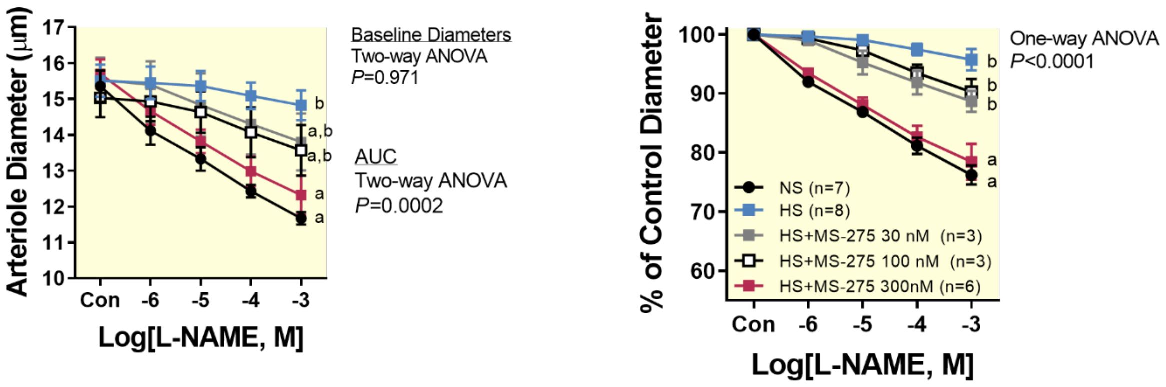


**Supplemental Figure 1.** L-NAME constricts afferent arterioles from normal salt (NS) fed rats in a concentration dependent manner. High salt diet (HS) significantly blunts constriction to L-NAME. MS-275 (300 nM) restores constriction to L-NAME in arterioles from (HS). Lower concentrations of L-NAME (30 nM and 100 nM) do not restore constriction to L-NAME back to NS levels. All arterioles were perfused with blood from a rat on a matching diet. Arteriolar diameters are presented as measured (left panel) and normalized to baseline (right panel). Data are plotted as mean±SE. *a* and *b* notate groups with statistically different areas under the curve (AUC). Groups noted *a,b* are not different than either groups labeled *a* or *b*. Differences in baseline diameters and AUC were assessed by two-way ANOVA and Holms-Sidak post hoc test.

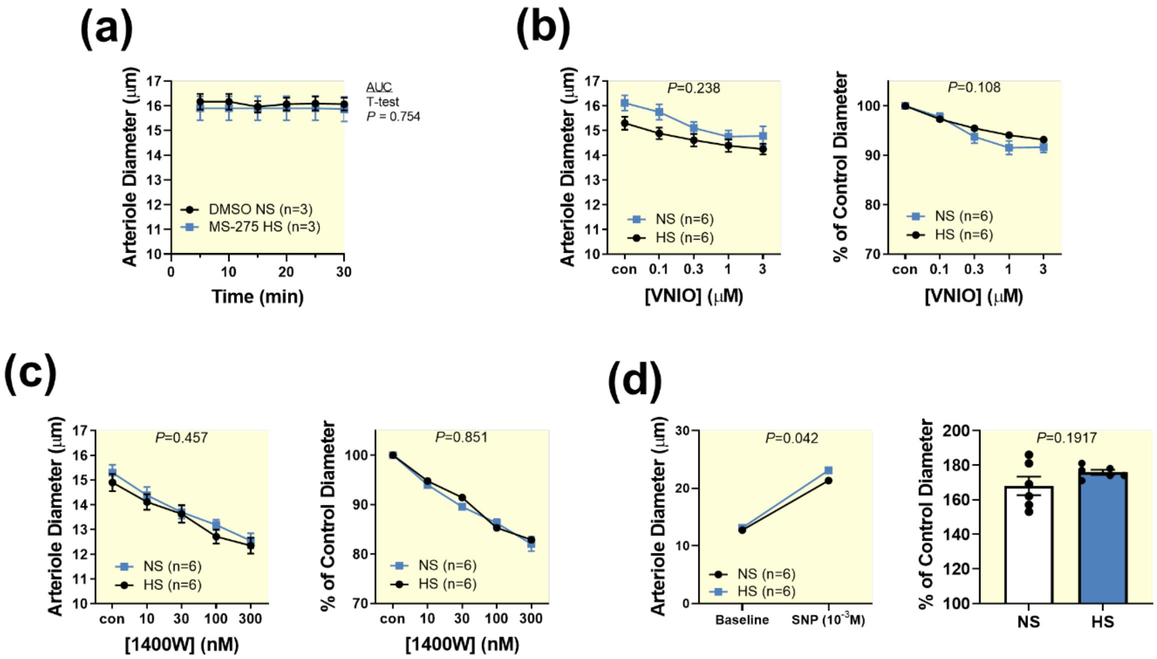


**Supplemental Figure 2. Afferent arteriole diameters for MS-275 intervention and reactivity to VNIO, 1400W, and sodium nitroprusside.** (**a**) DMSO (0.1% vol/vol) and MS-275 (300 nM) do not induce changes in afferent arteriole diameter. (**b-d**) Afferent arterioles from NS and HS fed rats react similarly to (**b**) the NOS1 inhibitor VNIO and (**c**) the NOS2 inhibitor 1400W. Area under the curve compared with unpaired, two-tailed Student's t-test. (**d**) Arterioles from HS fed rats show no impairment to sodium nitroprusside (SNP) dilation. HS fed rats showed slightly higher arteriolar diameters in response to SNP, but this difference was not observed when diameters were normalized to baseline diameter. All arterioles were perfused with blood from rats on a matching diet. All data are plotted as mean±SE. Comparisons were analyzed with unpaired, two-tailed Student's t-test

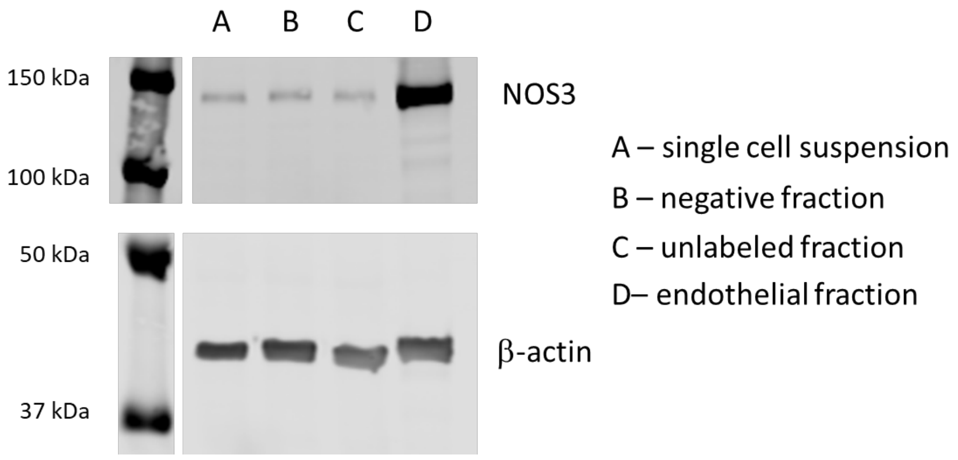


**Supplemental Figure 3. Endothelial Cell Isolation.** The four fractions generated during endothelial cell isolation were analyzed for NOS3 abundance. Fraction A: kidney single cell suspension; Fraction B: non-endothelial cell fraction which is positive for the negative selection markers; Fraction C: unlabeled cells; Fraction D: endothelial cells. The endothelial fraction contains much more NOS3 than the other fractions as expected.

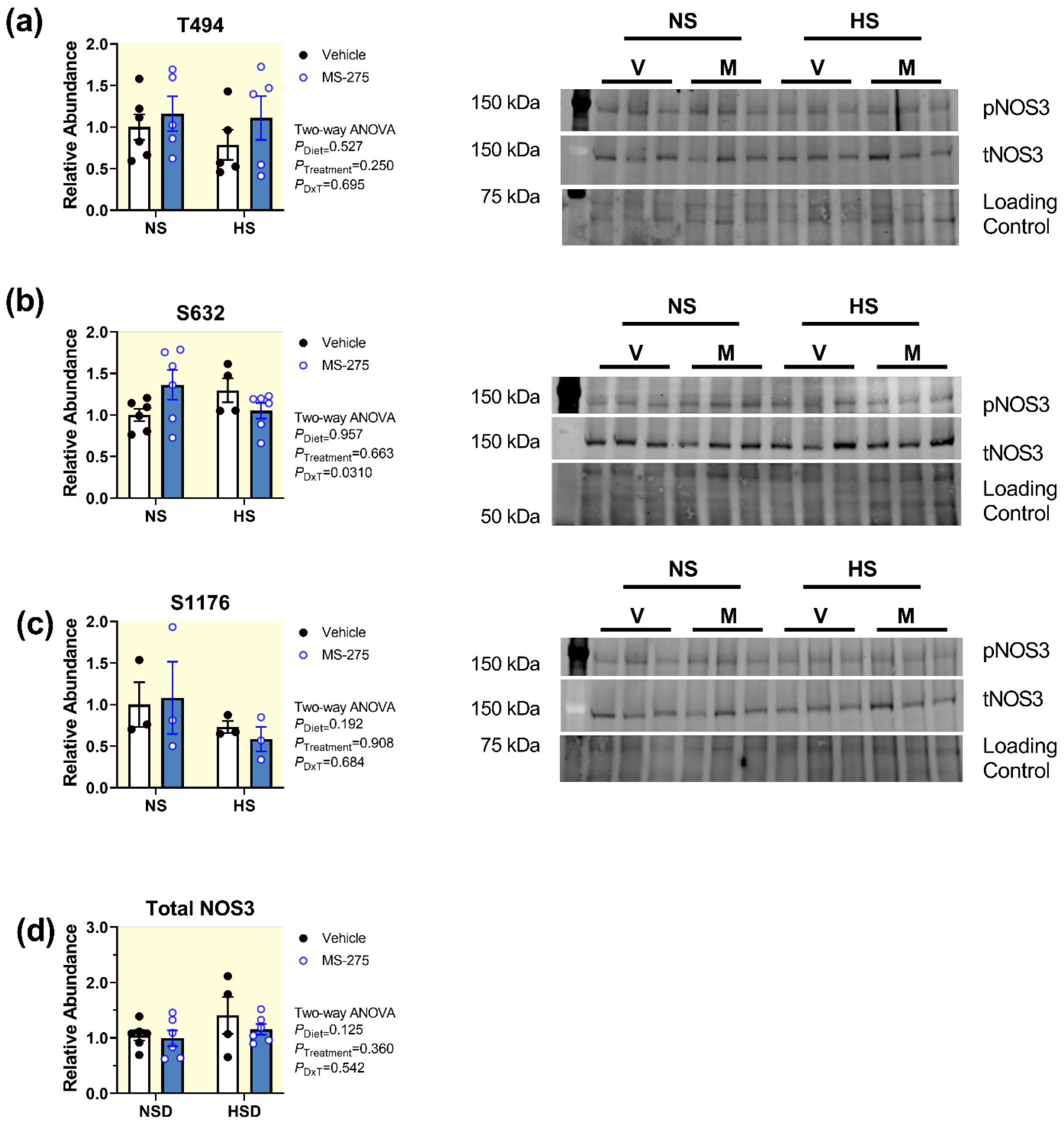


**Supplemental Figure 4. NOS3 phosphorylation in kidney vessels.** NOS3 phosphorylation at (**a**) T494, (**b**) S632, or (**c**) S1176 and (**d**) total NOS3 were unaffected by HS and HDAC1 inhibition with 300 nM MS-275. Differences were assessed by two-way ANOVA and Sidak post hoc test.
